## Supplemental Figures and Table for "CaBP1 and 2 enable sustained Ca_V_1.3 calcium currents and synaptic transmission in inner hair cells"

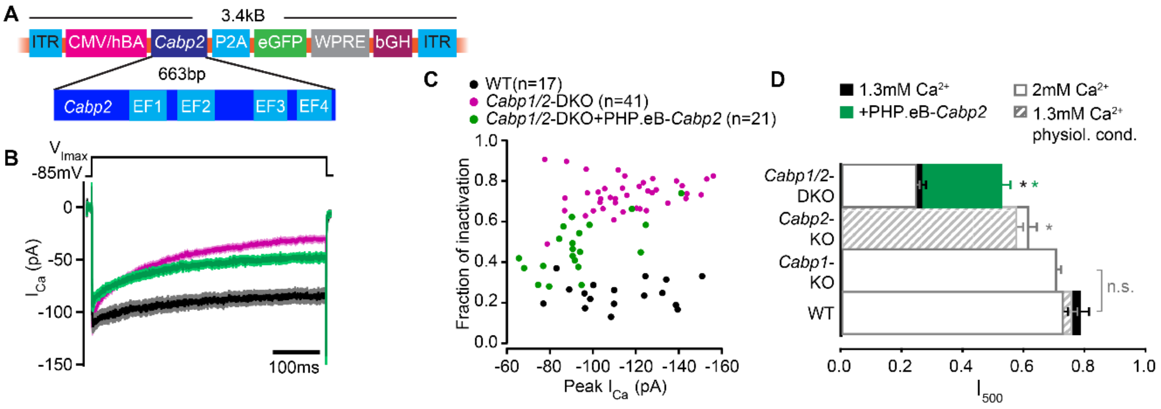


**Fig. S1. Partial rescue of $Ca_V1.3$ channel inactivation properties upon transgenic expression of CaBP2**. **(A)** The viral vector contains a hCMV/HBA (human cytomegalovirus immediate early enhancer, human beta actin promotor) hybrid promotor to drive strong expression of CaBP2 and a P2A sequence for bicistronic expression of the reporter gene *eGFP*. The Woodchuck Hepatitis Virus Posttranslational Regulatory Element (WPRE) and the bovine growth hormone (bGH) poly-adenylation sequence were included in the construct to enhance transcription and improve the stability of the transcript. **(B)** Absolute $Ca^{2+}$-currents after a 500-ms depolarization step to the maximum current potential in the apical IHCs of WT and *Cabp1/2*-DKO animals. **(C)** Peak calcium current and the corresponding fraction of inactivation after a 500-ms depolarization to the maximum current potential. Note no correlation between peak calcium current and fraction of inactivation, as is typically observed for large enough current amplitudes. **(D)** The fraction of the remaining $Ca^{2+}$-current after a 500-ms depolarization step is depicted for different mouse models and conditions. The histogram combines the results of this study (*Cabp1/2*-DKO; 1.3 mM [$Ca^{2+}$]: *n*=41, 2 mM [$Ca^{2+}$]: *n*=13) with previously published data on CaBP1[12] (*n*=20), CaBP2 acquired at physiological conditions [21] (*n*=14) and at room temperature[13] (*n*=14). The WT data in 2 mM extracellular [$Ca^{2+}$] (*white bars*) combines pooled data from the current study and controls as obtained previously[12] (*n*=17). For WT data acquired at RT and 1.3 mM [$Ca^{2+}$] *n*=17 and for physiological conditions [21] *n*=8. Note that the recordings from *Cabp2*-KO animals were acquired at the lower apex to mid-cochlear tonotopic positions as the phenotype in the apex (low-frequency positions investigated otherwise) is mild [13]. Asterisks represent statistical significance as compared to the corresponding controls (the results taken from the afore mentioned publications and current analysis).

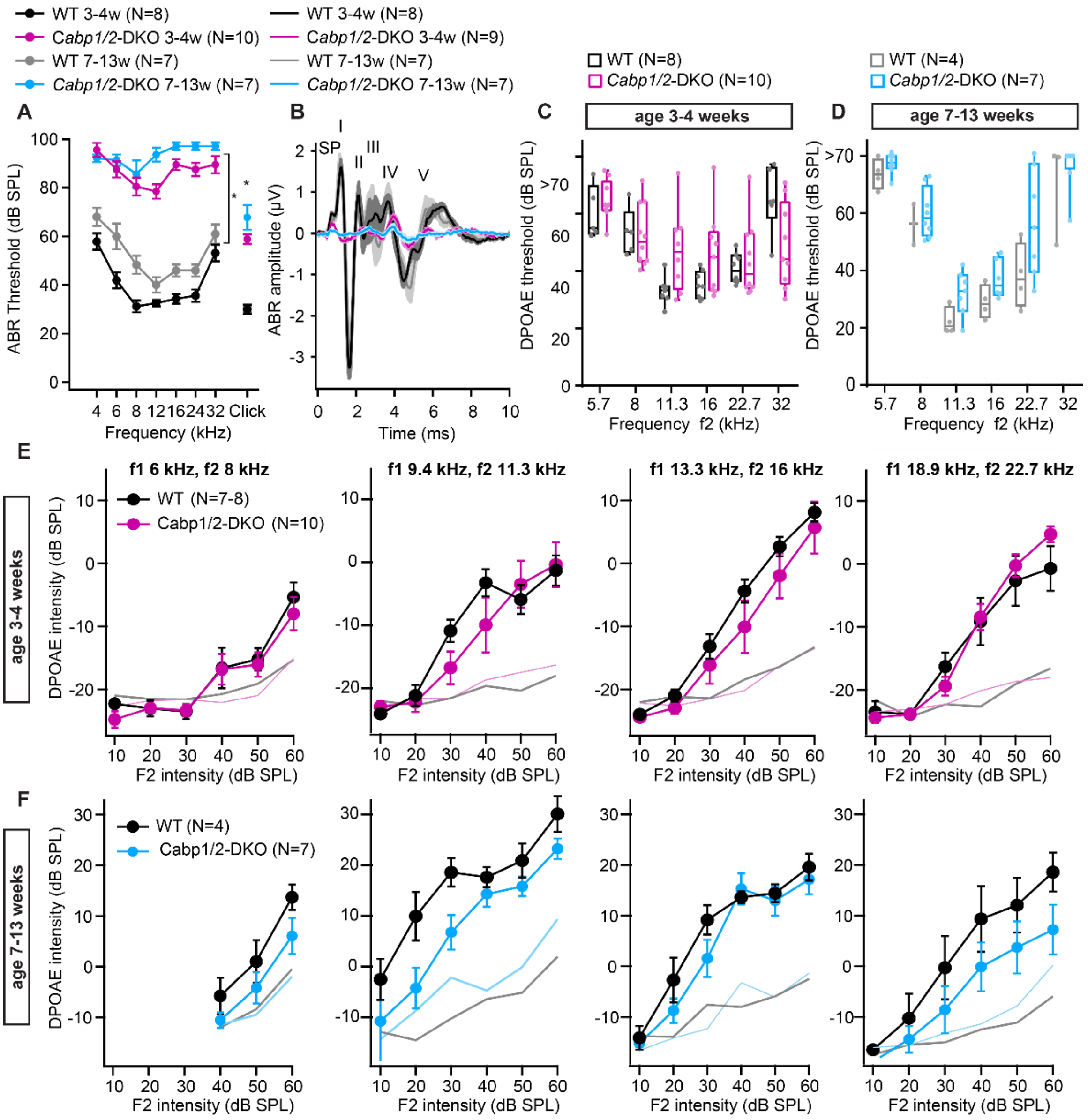


**Fig. S2. ABR amplitudes are strongly decreased while DPOAE are mostly preserved in the *Cabp1/2*-DKO animals. (A)** ABR thresholds to tone burst and click stimulation show an age-progressive pantonal severe hearing loss in *Cabp1/2*-DKO mice (asterisks: Šidak's multicomparison tests for tone burst, Student's *t* tests for clicks; $p \leq 0.0001$). When thresholds exceeded 90 dB for the tone burst stimulation, they were assigned a value of 100 dB for calculation of the mean and SEM. **(B)** Grand averages of ABR waveforms to 80-dB click stimuli presented at a rate of 20 Hz show a strong decrease in ABR amplitudes in *Cabp1/2*-DKO mice at 3-4 weeks of age. At age 7-13 weeks there are only minimal responses remaining, which are dominated by a delayed wave IV. The ABR wave peaks are labeled by roman numbers. Wave I is preceded by the summating potential (SP). **(C-D)** An increase in DPOAE thresholds was observed in some *Cabp1/2*-DKO mice, and was significant at the age of 7-13 weeks ($p < 0.002$, 2-way ANOVA). However, note that even at 7-13 weeks of age some *Cabp1/2*-DKO mutants retained seemingly normal DPOAE thresholds. **(E-F)** DPOAE amplitude growth functions for different primary tone frequency combinations at the age of 3-4 weeks **(E)** and 7-13 weeks **(F)**. The two datasets (E and F) were acquired at different setups calibrated with different artificial mouse ear couplers, resulting in different absolute amplitudes. Light colored lines depict average noise floor level.

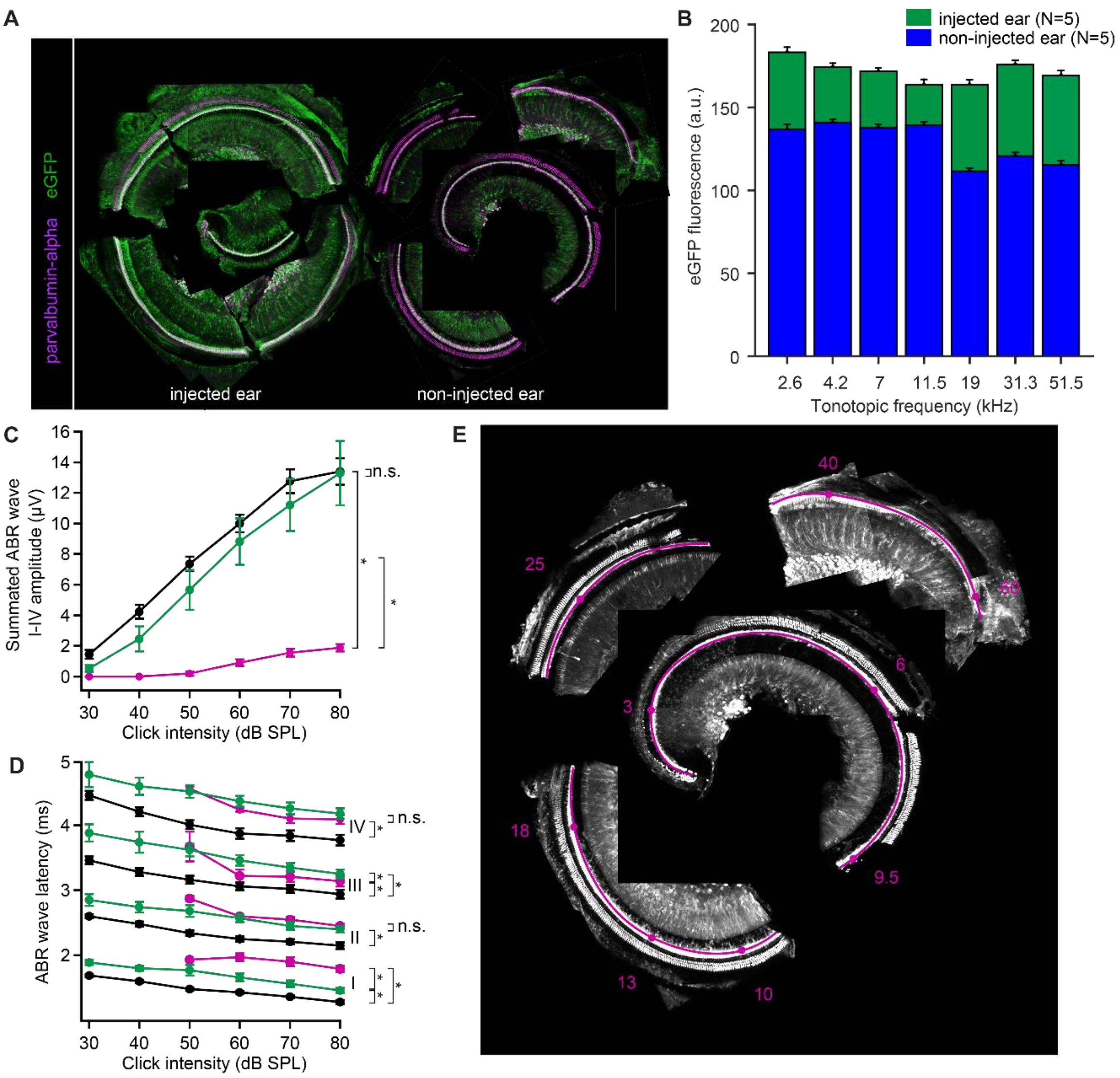


**Fig. S3. Efficient transgene-expression of *Cabp2* recovers cumulative ABR amplitude responses and ABR wave I latencies in *Cabp1/2*-DKO mice. (A)** Representative images of the organ of Corti of the cochlea immunostained with antibodies against parvalbumin-alpha as a hair cell marker and eGFP. Note a strong eGFP immunofluorescence detected along the entire tonotopic length of the injected and, to a lesser degree, contralateral non-injected cochlea. Scale bar: 20 µm. **(B)** eGFP immunofluorescence at each tonotopic position in the injected and contralateral non-injected ears. In both ears, but particularly on the contralateral side, lower eGFP expression in IHCs was observed towards the cochlear base. **(C)** A strong recovery of the summated ABR waves I-IV (Tukey's multicomparison test; $p < 0.0001$ for DKO vs WT or vs injected DKO) upon *Cabp2*-transgene expression. **(D)** ABR waves in the *Cabp1/2*-DKO animals are delayed. Intracochlear injection of PHP.eB-*Cabp2* leads to partial recovery of the wave I latency (for statistical comparisons, see Table S1). **(E)** A representative image of a dissected organ of Corti and the corresponding frequency map (tonotopic frequencies in kHz, *red*).

| Adjusted *p*-values | | *Cabp2*-injected DKO animals vs control DKO animals | DKO vs WT animals | *Cabp2*-injected DKO animals vs WT animals |
|---|---|---|---|---|
| **ABR thresholds** | 6 kHz tone burst | **** | **** | 0.4 |
| | 12 kHz tone burst | **** | **** | 0.002 |
| | 24 kHz tone burst | **** | **** | **** |
| | 20-Hz click | **** | **** | < 0.03 |
| **ABR wave latencies** | Wave I | **** | **** | **** |
| | Wave II | 0.1 | **** | **** |
| | Wave III | < 0.03 | *** | **** |
| | Wave IV | 0.1 | **** | **** |
| | Wave V | 0.9 | **** | **** |

**Table S1. Statistical analysis of ABR thresholds and ABR wave I-V latencies**. The table lists the adjusted *p*-values of the Tukey's multicomparisons tests upon one-way (20-Hz clicks) or two-way ANOVA (otherwise) for the ABR thresholds and the click-evoked ABR wave I-V latencies. Very low *p*-values are represented by asterisks: $p < 0.001$ (***), $p < 0.0001$ (****).
